# Sustained hovering, head stabilization and vision through the water surface in the Pied kingfisher (*Ceryle rudis*)

**DOI:** 10.1101/409201

**Authors:** Gadi Katzir, Dotan Berman, Moshe Nathan, Daniel Weihs

## Abstract

Pied kingfishers (*Ceryle rudis*) capture fish by plunge diving from hovering that may last several minutes. Hovering is the most energy-consuming mode of flight and depends on active wing flapping and facing headwind. The power for hovering is mass dependent increasing as the cube of the size, while aerodynamic forces increase only quadratically with size. Consequently, birds above a certain body mass can hover only with headwind and for very short durations. Hummingbirds are referred to as the only birds capable of hovering without wind (sustained hovering) due to their small size (ca. 2-20 gr), high wing-beat frequency and unique anatomy.

We studied the hovering characteristics of pied kingfishers in relation to wind and sun orientation, in 139 hovers. Furthermore, plunge diving necessitates the coping with the visual effects of light at the air/water interface. The kingfishers oriented their body axis towards the wind more than towards the sun. Hovering in little or no wind was common. With increased wind speed (a) orientation precision increased, (b) wing beat amplitude did not change, (c) wing beat frequency decreased and (d) body tilt became more horizontal. The head was highly stabilized and with orientations that indicated monocular viewing of prey.

We conclude that pied kingfishers achieve sustained hovering. This is despite their being significantly heavier than the theoretical maximum and showing ordinary kinematics and morphology. Head stabilization is a means of aiding viewing of submerged prey across the interface.

## Introduction

Many fish eating (piscivorous) birds plunge-dive into water from perches, (e.g., kingfishers) or from a flapping flight, to capture fish. Of the latter, most plunge dive from a direct flight (e.g., brown pelicans, *Pelecanus occidentalis*, gannets, *Sula* spp. and *Morus* spp., terns, *Sterna* spp.), while only a handful (e.g., Osprey, *Pandion haliaetos*) plunge dive from a hovering flight, i.e., from a relatively stable position in the air, maintained by wing motion (Cramp et al., 1977).

Of the ca. 100 species of kingfishers (Aves, Alcedinidae), the majority prey predominately on terrestrial prey while only ca. 20 species prey predominately on fish. In both terrestrial and aquatic habitats the kingfishers employ “perch-diving” as a common mode to capture prey (Fry & Fry, 1992). A foraging pattern that is distinctly different is observed in the piscivorous pied kingfisher (*Ceryle rudis*, Cerylinae; Reyer & Westerterp, 1985; Reyer et al., 1988). Pied kingfishers (PKF) typically forage using flapping flight with intermittent hovering episodes, during which position is unchanged for a given duration and from which they plunge dive (Fig. 1; Cramp et al., 1977; Fry & Fry, 1992). Hover duration ranges between several seconds and several minutes and hover heights – from less than one to more than eight meters above the water.

**Fig 1.**
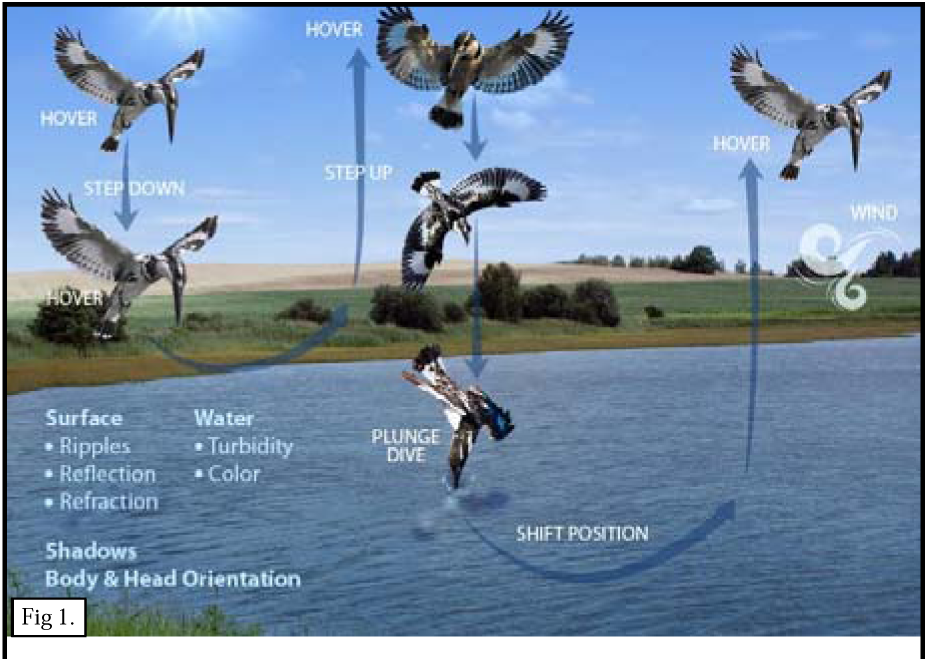
A schematic foraging path of a pied kingfisher. A hovering episode terminated if the kingfisher either (i) plunge-dived, or (ii) made a sharp vertical or horizontal change of position or(iii) landed, or (iv) left the area altogether.

The use of lengthy hovers is uncommon in birds and raises questions especially as to their capacity for sustained hovering – i.e., keeping a fixed spatial position without aid of wind. The forces required for hovering are provided by wing flapping and by utilizing wind, when available (Videler et al., 1983; Daan et al., 1990; Pennycuick, 1990; Pennycuick, 2008). The power required for hovering is mass dependent, increasing as the cube of the size, while the aerodynamic forces increase only quadratically with size, being dependent on wing dimensions. Sustained hovering is the most energy-consuming mode of flight, being ca. 4-5 times more energy demanding than regular flapping flight, because the weight of the bird has to be fully supported by its muscle power.Consequently, for birds above a certain body mass, hovering is very much wind dependent and birds such as kestrels, short toed eagles or osprey can hover in still air only for several seconds, yet perform significantly longer hovers in the wind (Grubb, 1977a,b; Machmer & Ydenberg, 1990). The contribution of wind to lift, however, becomes negative at high wind velocities, due to the increased drag and the destabilization by wind bursts.

Sustained hovering is mostly associated with hummingbirds (Trochilidae). This group shows extreme morphological, anatomical and physiological adaptations (Altshuler & Dudley, 2002; Tobalske, 2010) reaching peak aerodynamic performances that exceed present technical capabilities (Videler et al., 1983; Pennycuick, 2008). The notion prevailing in the literature is that hummingbirds are the only birds capable of sustained hovering (Chai & Millard, 1997), due to their small size, high wing-beat frequency and anatomy (Tobalske, 2010). We show that the much heavier pied kingfishers are capable of sustained hovering.

Plunge diving for fish requires coping with multiple visual effects of light at the air/water interface (Katzir & Intrator, 1987; Labinger et al., 1991). Wind is a major factor affecting vision through creating of water surface ripples and determining ripple direction, frequency and amplitude. Ripples and wavelets constantly change patterns of reflected and refracted sunlight and skylight that, in turn, affect image formation. We posit that a hovering PKF will attempt to place itself, and to orientate, so as to maximize the quality of the information gained on its underwater prey prior to the plunge dive that, once begun, is ballistic. The positioning is determined by the interaction between aerodynamic needs to produce lift and decease drag, as well as by the visual needs to reduce the effects of light refraction and reflection and the detection by the prey. Thus, for example, wind direction will result in a given 3D body orientation while concurrent visual requirements will result in a different 3D orientation of the head (Fig.1&10).

To answer the hypotheses above, we conducted field observations and discuss the results in the light of the potential pros and cons of hovering and of head stabilization.

## Materials & Methods

Field observations were conducted at the fisheries of Kibbutz HaMa’apil, Israel (Lat. 32.367348 Lon. 34.975555) in an area comprising 15 rectangular fish-ponds (width range 30-47m, length range 65-97m). Pied kingfisher foraging in the ponds was frequent and the surrounding low-lying vegetation (< 50cm) allowed direct observations over long distances. We conducted observations from December 2014 to July 2015 and the data analyzed was selected from 50 hours of observations performed over 13 days, between 06:00 and 20:00 so as to cover a range of wind, temperature and sun conditions.

A foraging sequence comprised one or more forward flight episodes interspersed with hovering flight episodes. Hovering episodes are defined as periods in which the bird moved less than one body length horizontally or vertically over many seconds. A sequence was analyzed from first sighting of a PKF to its leaving of the pond area. Most plunge dives commenced from stationary hovers (Douthwaite, 1976; Cramp et al., 1977; Reyer et al., 1988).

Hovering episodes were video recorded using a Canon© Powershot SX-50 camera on a tripod ca. 2.0m above theewater surfac e, with an optical zoom of 50X and filming at 24, 120 or 240 fps. A PKF was selected as a focal individual if it flew within ca. 50m of the observer. It was first tracked freely with the camera and once a hovering episode began, the camera was locked and the lens zoomed in on the subject. Overall, 375 sequences were acquired. Of these, 139 were selected as suitable for analysis in terms of duration and image quality. Video analysis was performed using a semi-automatic / manual tracking software developed by SIPL of the Technion and by “Kinovea©” software (version 0.8.15).

### Wind and sun

We recorded wind velocity immediately following the video recording of each foraging sequence. Measurements were performed using a handheld digital anemometer (Skywatch-Xplorer1), ca. 3m above the ground and ca. 4m above water surface. Wind direction was estimated from the compass direction of maximum wind velocity. Sun azimuth and elevation at each foraging bout was obtained from a web-available chart (http://www.sunearthtools.com/).

### Wing beat amplitude and frequency

For wing beat amplitude, we analyzed 33 hovering episodes. For the proximal wing, wingtip peak positions (highest / lowest) were marked in up to 11 consecutive cycles, and the vertical distance (in pixels) between any two consecutive peaks was measured. We used published measurements of PKFs (wingspan 46cm, wing length 19 cm; Cramp et al., 1977) to compute the angular displacement from stroke amplitude and the wing length. For wing beat frequency, we analyzed one sequence from each of 42 again randomly chosen episodes. We marked a frame at the very beginning of the first clearly measurable sequence and recorded the consecutive (up to a maximum of 10) wing beat cycles. Wing beat frequency was calculated from the number of frames required to complete a full cycle, at filming rates of 24, 120 and 240 frames per second (fps).

### Body orientation and tilt

To determine body orientation, (i.e., azimuth, in the horizontal plane) and tilt (in the vertical: see Fig.2) we selected 35 sequences in which the sagittal plane of the kingfisher as approximately perpendicular to the camera’s optical axis. From each sequence we selected a single episode and the orientation of the body (sagittal plane) was determined as follows: A single frame was captured from the beginning, middle and end of each selected episode. The body orientation relative to the camera was determined by eye, based on the shape of the body and the wing overlap, as viewed from the direction of the camera (Figs. 2 & S1). Body orientation was determined using rough blocks of ca. 22°, with 0° defined as viewing the PKF at right angles with its bill to the right. The absolute body orientation was calculated from the known azimuth of the camera’s optical axis. The orientation of the body relative to the wind and to the sun was calculated by subtracting the wind or sun values from body values.

**Fig. 2.**
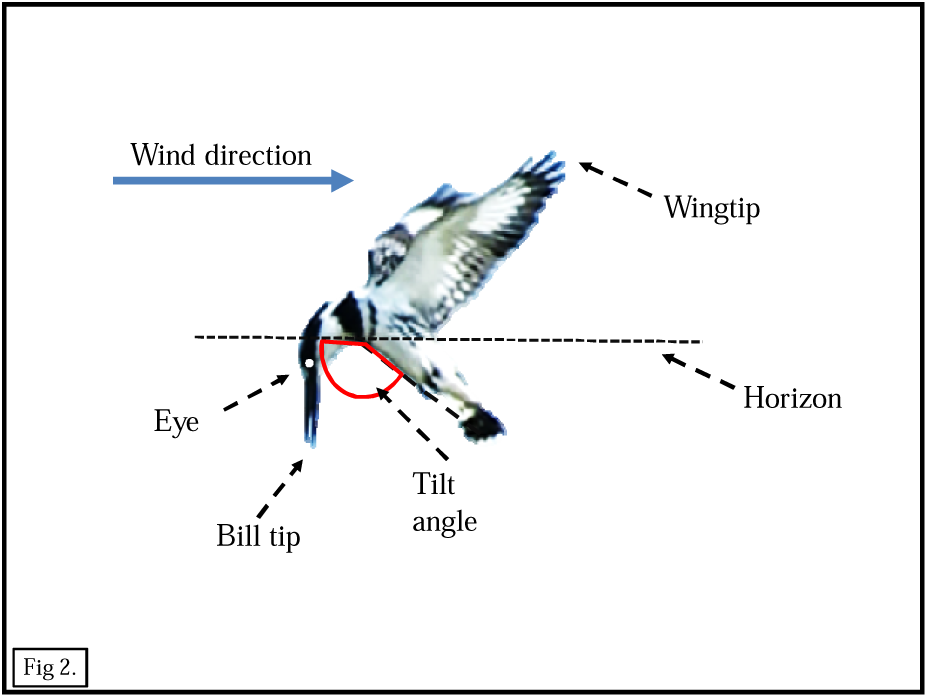
Parameters used in this study.

In hovering, the body underwent periodic changes in tilt, corresponding with wings’ down strokes. From 29 episodes, selected each from a different sequence, the orientation was analyzed for up to 10 consecutive cycles in which there was no apparent displacement of the center of gravity. Body tilt was determined from the angle of the imaginary line connecting the tip of the tail and the neck, with the horizon, (Figs. 2 & S1). The frames sampled were from points at which the body was closest to vertical. The absolute body tilt was calculated from the known azimuth of the camera’s optical axis as above. For each sequence sampled, the average of the body tilt angles was calculated.

### Movements of the head and body

For the analysis of movements and stabilization only sequences in which the sagittal plane of the PKF was approximately perpendicular to the camera’s optical axis were selected. Analyses were based on three points on the PKF’s body: The eye, the breast (approximately the rostral point of the sternum) and the proximal wingtip (Fig. 2). For six episodes we analyzed the change of vertical position of all three points while for the eye, horizontal change was also tested. For each episode, up to 10 wing beat cycles where sampled, from recordings taken at 120 fps, and the number of frames analyzed ranged between 90-240. For each of the points, the x-y coordinates of the first reading was taken as the origin (0,0) and subsequent readings were relative to it.

The angle of viewing of the PKF by the camera, in different hovering episodes, varied as a function of the relative 3D positions of the camera and the bird at the onset of each episode. To correct for these differences we used a photographic analysis of a mounted PKF. The PKF was secured horizontally on a stand, with its wings half open and its bill pointing down (Fig. S1). The origin (0,0) was the point in which the optical axis of the camera was horizontal, at the level of the PKF breast, and perpendicular to the sagittal plane of its body, at its center. Subsequent photographs were taken of the PKF from the predetermined intersection nodes of latitude lines (ventrally, 0°, - 30°, −60°) and longitude lines (rostral, 0°, +30°, +60°; caudal, 0°, −30°, −60°). From the real dimensions of the PKF and the photographic dimensions we calculated a correction factor for each viewing angle, using Kinovea (tm). The absolute tilt angle and bill length were taken from the images that were perpendicular to the bird’s sagittal plane.

The orientation of the PKF was analyzed using circular statistics of non-parametric angular-angular correlation (Zar, 1996). All other statistical tests were performed using SPSS (version 21). AIC ranking was used to determine models of best fit. Weather conditions during the research period ranged from hot, sunny days with little or no wind to stormy days with winds reaching 18 m/sec.

## Results

### Flight patterns in foraging

The kingfishers captured fish mostly by plunge diving from hovering flight or, infrequently, by diving from a perch. A typical foraging bout comprised episodes of level flight and episodes of stationary hovering (Fig. 1). The durations of individual hovering episodes ranged from a few seconds to more than a minute and the durations of entire bouts, from entering a pond area to leaving, ranged between a few seconds and over two minutes. Typically, a PKF would fly several meters above the pond in a fast, level flight, at an average speed of 10-12m/sec. It would then halt abruptly and hover in midair, at heights ranging between 0.5-12m, mostly with its bill pointing down (overall relative to the horizon 76.4° ± 9.2°; mean±sd). Flight segments between hovering bouts frequently were of a shallow u-shape. Following a plunge-dive it would either resume hovering, or land on a perch or leave the area (Cramp et al., 1977; Douthwaite, 1976; Reyer et al., 1988).

### Body orientation

In each hovering episode, the body orientation of the PKFs in the horizontal plane remained virtually unchanged (variance ca. 0 over 35 episodes). Thus, once a position has been adopted, the PKF minimized positional changes.

Body orientation was significantly correlated with wind direction (Fig. 3; α=0.05, N=35) and there was a trend, although not reaching significance, of a decrease in relative body orientation with increased wind velocity (Fig. 5). In other words, the higher the wind velocity the higher the precision of facing it directly.

**Fig 3a:**
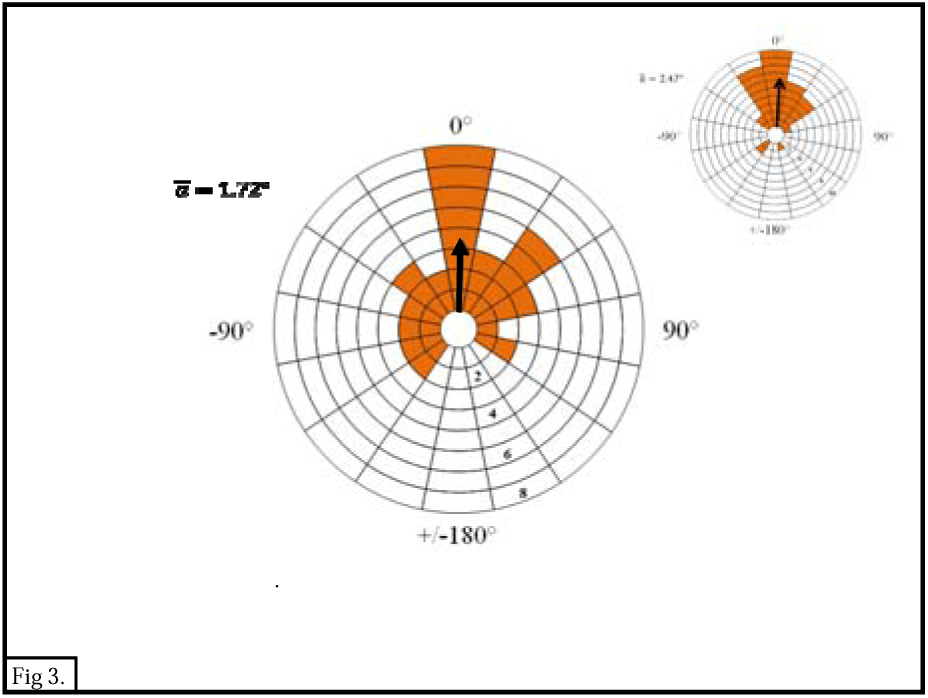
Body azimuth relative to sun azimuth. Vector diagram: Radius - highest number of observations in any sector (each sector - 22.5°). Azimuth 0° - facing the wind. Arrow - mean relative orientation (N=35 sequences). Fig. 3b. Difference between orientations to wind and to sun.

**Fig. 4.**
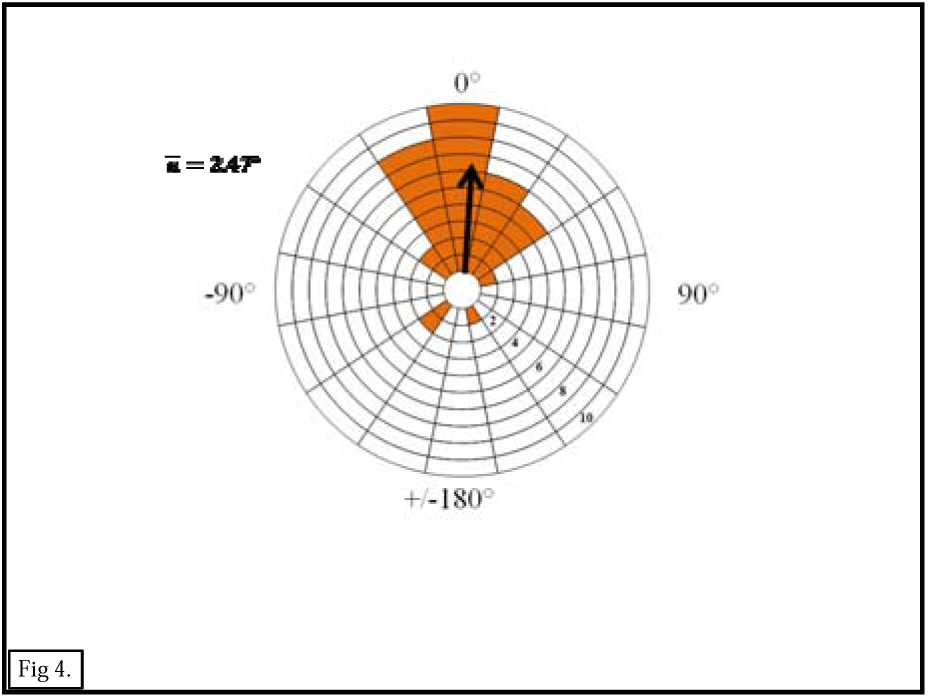
Body azimuth relative to wind azimuth. Vector diagram: Radius - highest number of observations in any sector (each sector - 22.5°). Azimuth 0° - facing the wind. Arrow - mean relative orientation (N=35 sequences).

**Fig 5:**
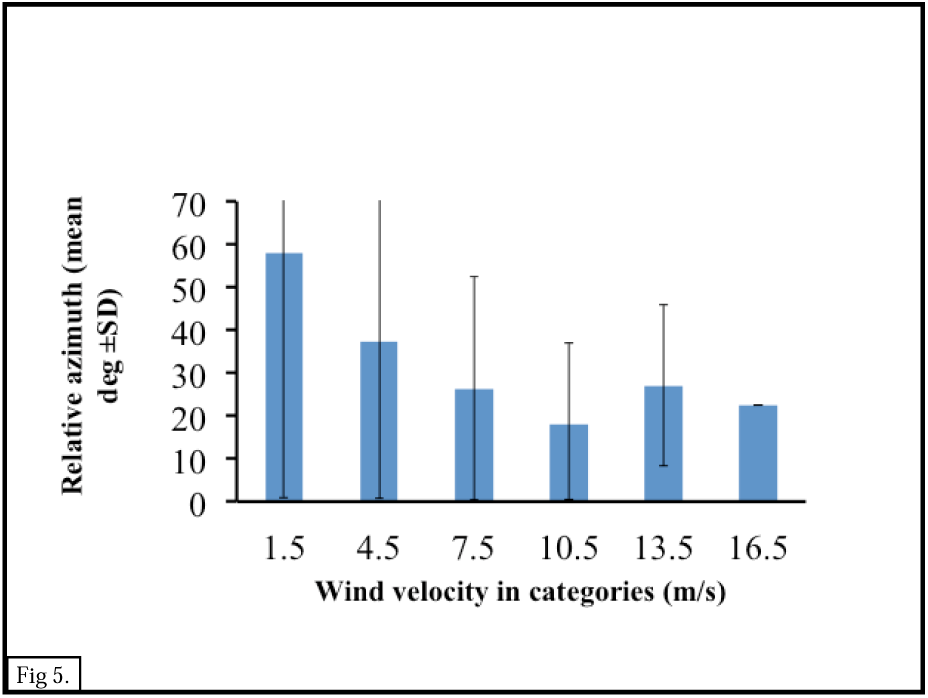
Body azimuth relative to wind velocity. Each column represents the mean ±SD of the relative body to wind azimuth in a given wind velocity category (N=35).

Body orientation was related to sun position, yet the correlation did not reach significance (Fig. 4; α>0.05, N=35).

### Motion of the wings, body and head

During a given hovering episode, the tilt angle of the body, the head-bill and the tail were roughly unchanged. The bill in most hovering episode pointed down (i.e., pitch) with the head partially rotated (Fig. 10) so that one eye gazed downwards. In the transition from a stationary hovering position to flight, the orientation of the head - bill first changed so that the bill first pointed in the direction of the ensuing flight, and the body followed.

**Fig. 6:**
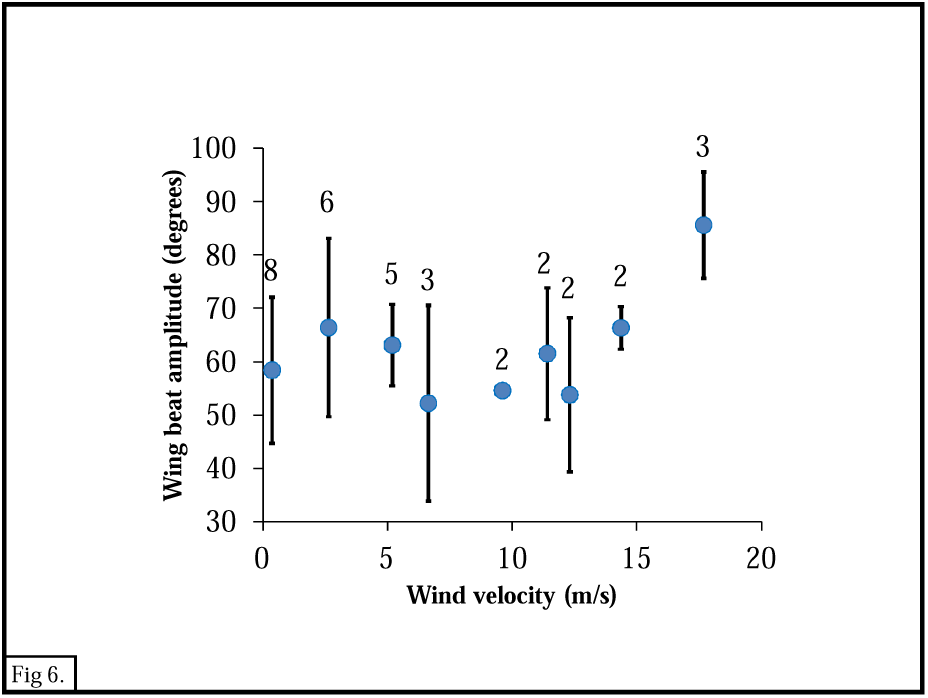
Wing beat amplitude relative to wind velocity. Each solid circle is the mean ±SD of between 3-10 wing beat cycles per sequence (N=33 sequences).

**Fig. 7:**
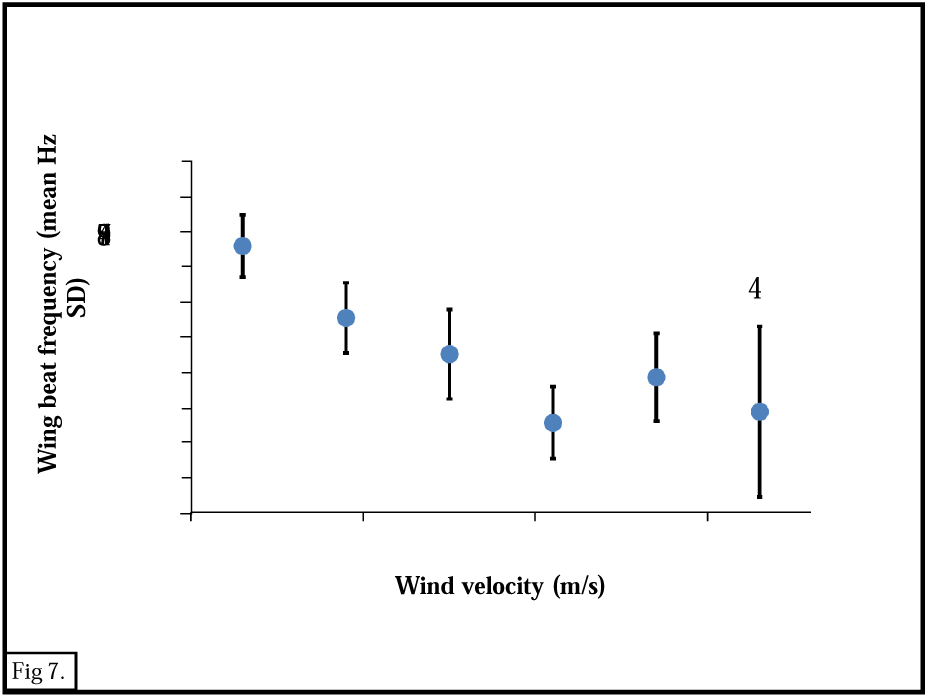
Wing beat frequency relative to wind velocity. Each solid circle is the mean ±SD of between 3-10 wing beat cycles per sequence (N=42 sequences).

**Fig. 8:**
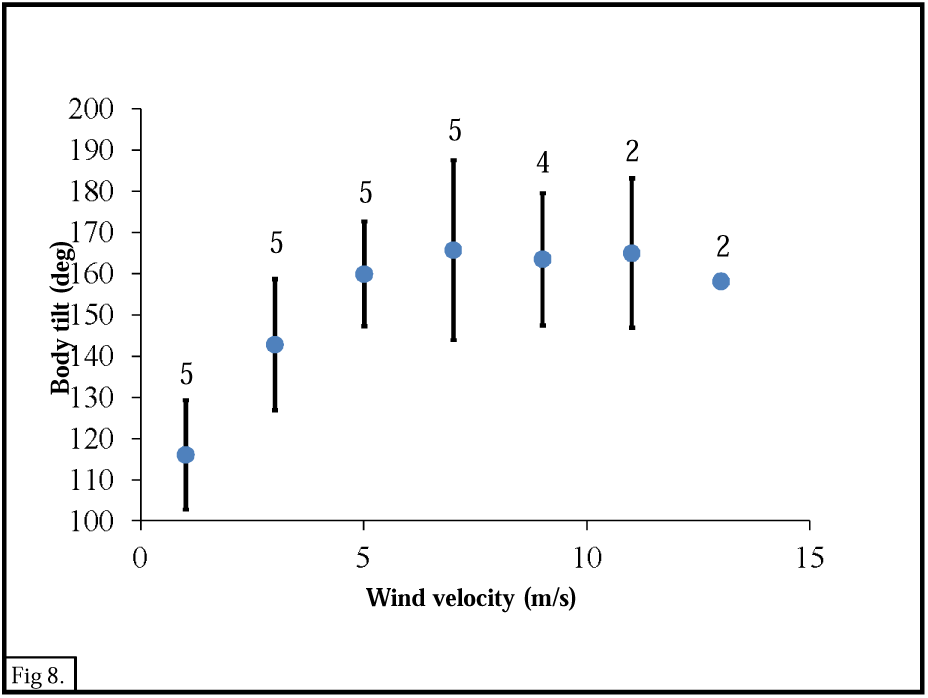
Body tilt relative to wind velocity (in categories). Each solid circle is the mean ± SD of the smallest angle of body tilt in each of 5-10 wing beat cycles (N=26 sequences).

**Fig. 9:**
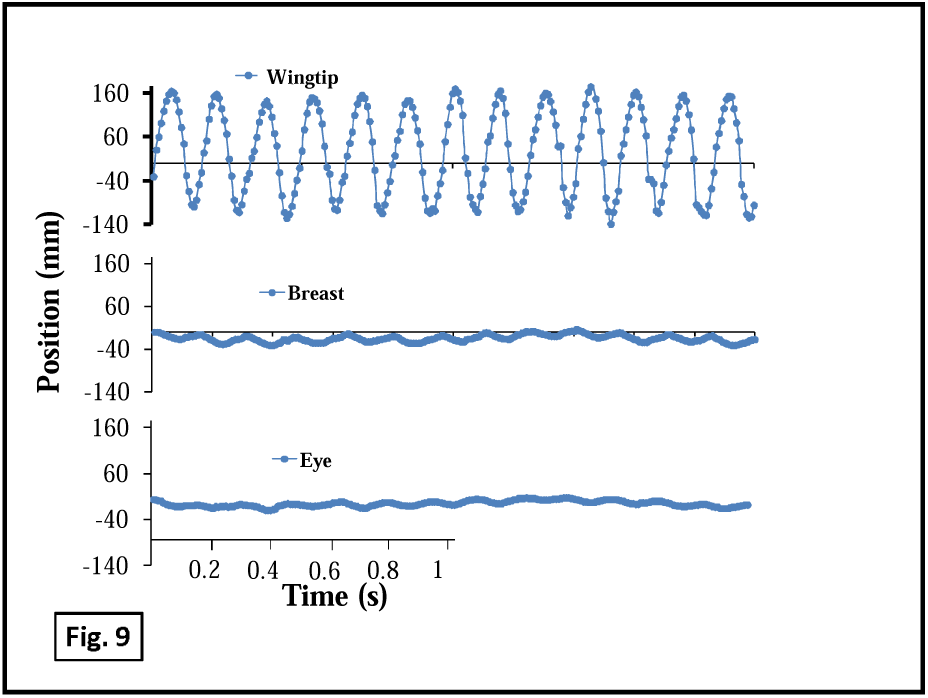
Stabilization of the head and body. Example of wingtip, breast and eye vertical position over seven wing beat cycles. Elements are drawn to the same scale. The smaller chart is in independent scales.

**Fig. 10.**
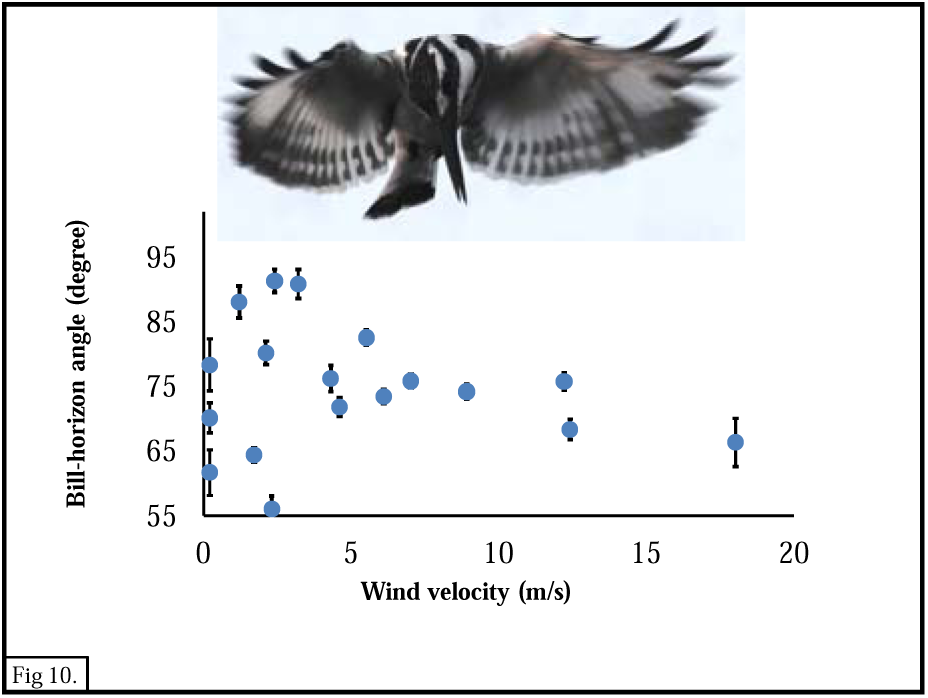
A. Head position relative to the torso and wings in a hover and a dive (inset). A clear roll and tilt are observed. B. Bill tip-eye angle relative to wind velocity. Each solid circle is the mean ± SD of bill tip-eye-horizon the (smaller) angle in each of 6-10 wing-beat cycles (N=18 sequences).

### Wing beat amplitude and frequency

Wing beat angular amplitude (range 9°-134°) was not correlated with wind velocity (Fig. 6) while wing beat frequency (range 5.4-9.5Hz) was significantly and negatively correlated with wind velocity (Fig. 7; N=42, R^2^= 0.725, P<0.05). In other words, the higher the wind velocity, the lower the wing beat frequency. Wing beat amplitude was minimal at wind velocity of ca. 7m/sec and wing beat frequency was minimal at ca. 12-14m/s (Figs 6 & 7).

During a hovering episode, the kingfishers’ body tilt underwent periodic changes with wing beat cycles, as the majority of the lift during the entire cycle is produced during the down stroke so that the body is moved upwards. The tilt serves to reduce the body drag during this upward motion of the cycle. The maximal vertical extent of body tilt was significantly correlated with wind velocity (Fig. 8) so that with increased wind velocity (up to ca. 12 m/s) the tilt became more horizontal. At wind velocities higher than 12 m/s, body tilt became more vertical again (N=29, R^2^=0.851, P=0.003).

### Stabilization of the head and body

The vertical displacements of the wingtips, body and head during a hover episode differed significantly, with the head (eye) undergoing the smallest displacements and the wingtip – the highest. In the example given (Fig. 9) the wingtip displacement (172.6 ± 72.7mm) were ca. two orders of magnitude greater than that of the breast or eye (respectively 8.4±4.6mm and 4.3± 2.7mm). Eye horizontal displacement was of similar magnitude (3-5.9mm). The periodicity of wingtip position differed from that of the eye and breast by 180° (Fig. 9).

## Discussion

Pied kingfishers are exceptional in making extensive use of hovering from which they plunge dive for fish, sometimes from heights well over 10m above the water (Cramp et al., 1977; Labinger et al., 1991; Fry & Fry, 1992). This falls within the general pattern of birds that hover near potential prey that mostly leads to a rapid strike, dive or plunge-dive (e.g., kestrel, *Falco tinnunculus*; short-toed eagle, *Circaetus gallicus*: osprey, *Pandion haeliaetus)*

Hovering is considerably more energy demanding than level flight so it is much less used. This is especially apparent in relation to hovering in still air (“sustained hovering”) a task that most birds are unable to perform as they cannot produce enough lift through muscle power (Videler et al., 1983) and must gain additional lift by flying into moderate wind (Videler, et al. 1983; Daan et al., 1990). The frequent and lengthy hovering episodes of kingfishers in still air prove their capacity for sustained hovering. This finding revises the prevailing view that sustained hovering is confined to extremely small birds. Previous theoretical and experimental approaches have led to the conclusion that maximal body size for sustained hovering is that of hummingbirds (5-10gr). Indeed they are regarded as “…the *only birds that can sustain hovering*…” (Pennycuick, 1978; Chai & Millard, 1997; Altshuler & Dudley, 2002; McNeill Alexander, 2005; Tobalske, 2007; Pennycuick, 2008;) due to “.. their small size, high wing beat frequency, … wing anatomy that enables them to.. supinate during upstroke so that they can generate lift on both up - and down-strokes …” (Chai & Millard, 1997; Tobalske et al., 2003; Tobalske, 2010; Goller & Altshuler, 2014). Pied kingfishers (ca. 80-100 gr) are considerably heavier than hummingbirds, their wing-beat frequencies are lower (ca. 7-10 Hz. Vs. 70-80Hz) and they do not exhibit morphological or motor specializations of the wings (e.g., unique aspect ratio or wing-load). Also, their flight kinematics seems most similar to non-hovering, flapping flight including wing-beat frequency and amplitude that are correlated with wind velocity and direction. Thus, it is not possible at this stage to point to specific aspects of the hovering flights that enable these capacities, unobserved in other species This leaves open questions of size limits for sustained hovering.

Hovering in the kingfishers was sensitive to wind and with increased wind velocity the birds orientated increasingly more closely into the wind, most probably to decrease the drag coefficient. As wind speed increases, the need to stay in a horizontally fixed position becomes more demanding, because drag increases quadratically with wind speed and so becomes more dominant. This turning point in the relative importance of drag is related to the volitional speed of the pied kingfisher forward flight. With increased wind velocity, body tilt became increasingly more horizontal, to reduce drag by reducing body frontal area. As wind velocity increased from zero to roughly 12 m/s, wing beat frequency decreased but wing-beat amplitude did not. We attribute this to the increasing contribution of wind energy to the production of the lift required to stay at a fixed altitude. At the higher wind velocities, hovering was infrequent and rather chaotic, as expected, because at these velocities every deviation due to local turbulence is of the order of the bird size. There seem to be two ranges of wind velocities, in which hovering have different characteristics, with the change occurring at about 12 m/s (Figs 3 and 4). These patterns are expected if the pied kingfishers were attempting to maximize lift while minimizing their own active contribution.

In hovering, the kingfishers kept their torso and their head fixed to different degrees. While the wings underwent displacements of up to ca. 120deg, the torso motion was reduced by an order of magnitude and the head movement was further reduced so that eye perturbation was of order 5mm. These results are similar to head stabilization of kestrels (Videler et al., 1983, pied kingfishers, unpublished) and herons on a moving perch (Katzir et al., 2001) While body tilt changed with wind speed, the orientation of the stabilized head was kept with the bill pointing downwards (Fig. 1, top center).

As to the pros and cons to hovering, one potential disadvantage is increased conspicuousness: To the fish, the kingfisher presents a high contrast silhouette against the sky mostly perpendicularly above, which is at the center of the fish Snell’s window, where optical distortions are minimal. Add to the attractiveness is wing motion. On the advantage side, hovering releases the kingfishers from the need to use vegetation to perch on, thus extending their foraging to several km offshore (Fry & Fry, 1992). Also, hovering provides a stable “springboard” that enables a rapid re-positioning in 3D to improve visual detection of prey under the visually complex air/water interface. Being perpendicular to the fish may reduce optical problems while dynamic re-positioning may reduce the problem of the rapid escape responses of fishes (Katzir & Camhi, 1993). So why is the pied kingfisher’s hovering so dominant among piscivorous birds? One factor may be the prey. Cichlids (Cichlidae), the most species-rich fish group, evolved in the African lakes and are a main prey of pied kingfishers. Many cichlids nest in shallow waters and the males keep a position above the substrate built nests making them conspicuous and vulnerable to avian predation. It may well be that pied kingfishers have evolved to cope with single, relatively large prey, requiring high precision. In comparison, species such as terns or gannets prey on fish in schools with large numbers of fish in motion, and where precise estimates can only be made at close range.

While hovering the kingfishers stabilize their eyes/head. Birds are known to stabilize their eyes extremely well even under marked perturbations, while flying or when walking. In the case of the PKF hovering can be performed w/o head stabilization (Videler et al., 1983; Troje & Frost, 2000; Warrick, 2002; McArthur & Dickman, 2011). Gaze stabilization of minimal duration is required to reduce image slippage on the retina and hence reduce blur (Land, 1999; Zeil et al., 2008). The sensory information for stabilization may be visual, such as optic flow (e.g., hummingbirds, Goller & Altshuler, 2014) or mechanical (inertial, vestibular, e.g., swans, Pete et al., 2015). As for using vision, there are several potential difficulties: Visually distinct “anchor points”, both beneath and lateral to the hovering bird, are relatively distant and most probably do not provide the precision provided by a nearby object such as a flower several cm from the eye of a hovering hummingbird. Moreover, the water surface provides dynamic patterns and unlikely to be used reliably. Thus a more probable source is the vestibular.

A clue to the constraints on the kingfishers’ vision may be found in the marked 3D tilt/roll orientation of head during the hovers (Figs. 1,10) that may stem from preferred gaze directions. In many birds, including kingfishers, each retina has a temporal fovea that “looks forward” and provides for close range binocular vision and a central fovea that “looks sideways” and provides for long distance, monocular, high resolution vision (Moore et al., 2015). Consequently, when faced with a distant target, such birds tilt their head to direct their monocular gaze at the target (Moroney & Pettigrew, 1987; Tucker, 2000). The pronounced head tilt / roll of the pied kingfisher allows monocular viewing of the prey while minimizing the aerodynamic negative effects of body orientation.

It may be useful to view a kingfisher as comprising two rigid units that interact, a larger torso and a smaller head, with three flexible units, the neck, wings and tail (Katzir et al., 2001). The body, wings and tail perform mainly the flight tasks and need to maximize lift, while affecting vision and the head orientates so as to maximize visual input yet also affects flight.

## Acknowledgements

We thank Kibbutz Ha-Maapil and the operators of the kibbutz fish ponds.

## Supplement

S1. Correction method. A set of images were made of a stuffed PKF. The photographs were from different angles that were calculated by positioning the camera at measured distance. from the object.

